# A new web resource to predict the impact of missense variants at protein interfaces using 3D structural data: Missense3D-PPI

**DOI:** 10.1101/2023.01.24.525222

**Authors:** Cecilia Pennica, Gordon Hanna, Suhail A Islam, Michael JE Sternberg, Alessia David

**Author notes:** joint last authors (Michael JE Sternberg, Alessia David).

## Abstract

In 2019, we released Missense3D which identifies stereochemical features that are disrupted by a missense variant, such as introducing a buried charge. Missense3D analyses the effect of a missense variant on a single structure and thus may fail to identify as damaging surface variants disrupting a protein interface i.e., a protein-protein interaction (PPI) site. Here we present Missense3D-PPI designed to predict missense variants at PPI interfaces.

Our development dataset comprised of 1,279 missense variants (pathogenic n=733, benign n=546) in 434 proteins and 545 experimental structures of PPI complexes. Benchmarking of Missense3D-PPI was performed after dividing the dataset in training (320 benign and 320 pathogenic variants) and testing (226 benign and 413 pathogenic). Structural features affecting PPI, such as disruption of interchain bonds and introduction of unbalanced charged interface residues, were analysed to assess the impact of the variant at PPI.

Missense3D-PPI’s performance was superior to that of Missense3D: sensitivity 42% versus 8% and accuracy 58% versus 40%, p=4.23×10^−16^ However, the specificity of Missense3D-PPI was slightly lower compared to Missense3D (84% versus 98%). On our dataset, Missense3D-PPI’s accuracy was superior to BeAtMuSiC (p=2.3×10^−5^), mCSM-PPI2 (p=3.2×10^−12^) and MutaBind2 (p=0.003).

Missense3D-PPI represents a valuable tool for predicting the structural effect of missense variants on biological protein networks and is available at the Missense3D web portal (http://missense3d.bc.ic.ac.uk/missense3d/indexppi.html).

## INTRODUCTION

Residues on the protein surface that are not directly involved in function generally are more tolerant to amino acid substitutions compared to residues affecting the buried core of a protein [1]. However, interface residues involved in protein-protein interaction (PPI), also known as interface residues, are an exception to this principle. As previously demonstrated by our group and others [1] [2], interface residues are enriched in disease-causing amino acid substitutions. The damaging effect of variants affecting protein interaction sites is difficult to predict and the majority of *in silico* variant prediction methods perform worst on variants located at the interface compared to the remaining protein surface or the buried interior protein area [3].

Genetic variants causing the disruption of protein interfaces are an important contributor to human disease [4] [5]. These variants generally preserve the folding and stability of the monomeric protein but may impair its function by impacting on the many biological processes which rely on protein interaction, such as trafficking and signalling. Identification and prediction of the effect of a variant on PPI requires knowledge of the residues forming a protein interface. In recent years there has been an increase in the availability of three-dimensional structures of protein complexes, both experimentally solved and obtained from protein docking and homology modelling [6]. These 3D coordinates are publicly available from databases, such as PDB [7], Interactome3D [8],GWYRE [9] and PrePPI [10]. Although at present the coverage of the protein interactome remains limited, we can expect an exponential increase in 3D coordinates of PPI complexes in the coming years as a result of the recent breakthrough in protein modelling achieved by AlphaFold and similar approaches which use deep learning [11] [12].

The availability of 3D coordinates allows us not only to predict the damaging effect of a variant, but also to understand the molecular mechanisms by which it affects protein structure/function. In 2019, we launched Missense3D [13], which predicts the effect of a variant on the folding and stability of a monomeric protein. However, Missense3D, similar to other algorithms such as HOPE [14] and SAAP [15], uses the 3D coordinates of a single protein chain, thus, potentially failing to identify the detrimental effect of variants located on the protein surface that may affect PPI. Such a damaging effect can only be predicted when the 3D coordinates of a protein complex are taken into account, an approach used by algorithms such as the energy-based programs BeAtMuSiC [16], MutaBind2 [17] and mCSM-PPI2 [18]. However, most of these *in silico* prediction tools have been trained on engineered protein variants deposited in databases, such as Skempi [19] and Protherm [20] and do not perform equally well when used on other datasets of variants [21] [22].

To date, the use of 3D structures to predict the effect of a variant remains relatively limited compared to the use of sequence conservation, thus calling for the development of new user-friendly algorithms that can be easily implemented to enhance variant prediction. We present Missense3D-PPI, a purely structure-based algorithm for the prioritization and characterization of missense variants occurring at protein-protein interfaces. Missense3D-PPI is available at the Missense3D web portal (http://missense3d.bc.ic.ac.uk/missense3d/indexppi.html).

## ALGORITHM

### Experimental structures and missense variants

Figure 1 presents the Missense3D-PPI pipeline. We extracted ~4 million human missense variants from our in-house Missense3D-DB database [23] which contains the phenotypic annotation of variants from ClinVar [24] and UniProt [25] and minor allele frequency (MAF) data from GnomAD [26].

**Figure 1.**
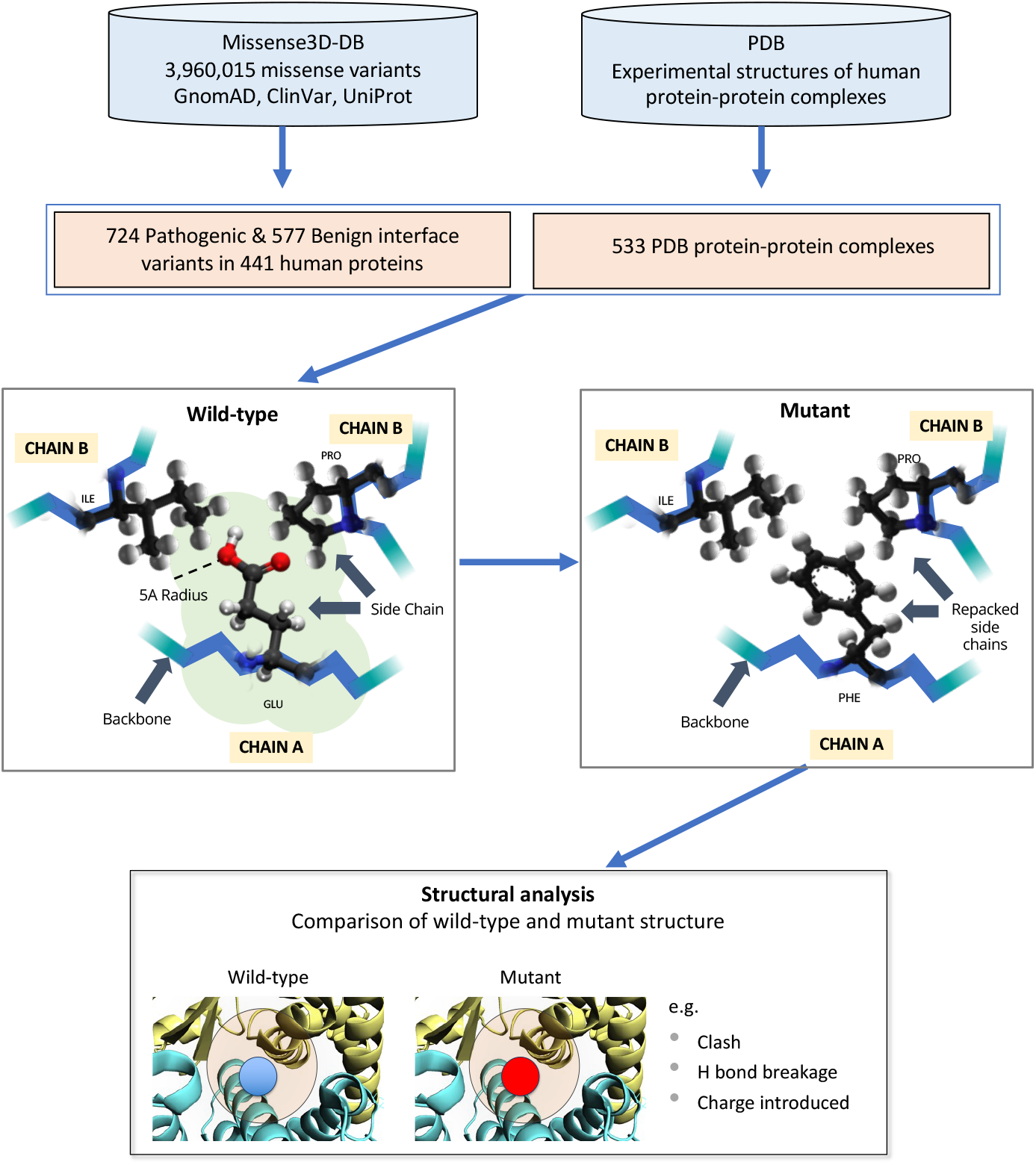
Missense3D-PPI pipeline used to analyse the structural effect of amino acid substitutions.

In order to identify missense variants occurring at a PPI site, we extracted 16,609 high resolution (≤ 2.5Å) X-ray crystal structures of human dimers and multimers from the Protein Data Bank (PDB) [27]. For each protein complex, we selected the experimental structure with the best resolution and without mutations in the protein interface. Interface residues were defined as any residue with a relative solvent accessibility (RSA) difference ≥5% between the monomeric and the protein complex structure calculated using an in-house program. Each interface residue was categorised as core, rim or support according to the change in RSA between the monomeric (RSA_monomer_) and complex (RSA_complex_) form:

- core residue: RSA_monomer_ ≥ 9% and RSA_complex_ < 9%; RSA_monomer_ - RSA_complex_ ≥ 5%
- rim residue: RSA_monomer_ ≥ 9% and RSA_complex_ ≥ 9%; RSA_monomer_ - RSA_complex_ ≥ 5%
- support residue: RSA_monomer_ < 9% and RSA_complex_ < 9%; RSA_monomer_ - RSA_complex_ ≥ 5%

We based our definition of core and rim according to our definition of change between buried and exposed status [13], but we acknowledge that several other definitions can be found in the literature. Only variants occurring at interface residues were retained and the final dataset comprised of 1,279 missense variants (pathogenic n=733, benign n=546) in 434 proteins and 545 PDB coordinates of PPI complexes. The benign dataset included variants with an annotation of “benign” from ClinVar or UniProt and variants with MAF>1% and no “pathogenic” annotation in ClinVar and/or UniProt. Human Leukocytes Antigen (HLA) proteins were excluded from the final dataset because of their highly polymorphic antigen binding amino acid residues [28]. Over 1,000 naturally occurring haemoglobin variants have been described and extensively studied clinically and in vitro [29]. In our dataset, haemoglobin variants annotated as “unstable” were also included in the “damaging” dataset.

### Missense3D-PPI pipeline and definition of damaging structural features

For each variant a mutant structure was generated using SCRWL [30], as described in Ittisoponpisan *et al*. [13]. Briefly, the mutant structure was generated by removing the side chain of the wild-type query residue and the side chain of all residues within 5Å distance from the query residue (defined by any pair of inter-residue atoms closer than 5Å). Subsequently, the side chain of the mutant residue was re-introduced and the structure repacked. The wild type and mutant structures were compared, and a variant was considered damaging if one of the structural features presented in Table S1, affecting two or more interchain residues was identified. The features “*Interface Gly, Tyr or Trp replaced”* - defined as the substitution of an interface amino acid from glycine, tyrosine or tryptophan to any residue - were introduced because of the functional importance of these residues within the protein interface [31] Moreover, similarly to Missense3D, in Missese3D-PPI 1Å was added to hydrogen bonds and salt bridges cut-off distances to account for the use of 3D models and structures with poor resolution [13].

### Evaluation of performance and benchmarking

The final dataset of 1,279 variants was split into a training and a testing set. To avoid an overlap of homologous proteins between the training and testing sets, homology between proteins was assessed using the HH-suite [32]. Clusters of homologous proteins were distributed to the training and testing set to obtain a similar ratio of variants in the two sets. The characteristics of the final datasets are described in Table S2.

The performance of Missense3D-PPI was calculated as per Missense3D algorithm [33]. Briefly, variants causing at least one damaging structural feature according to Missense3D-PPI were considered true positives if annotated as damaging, otherwise false positive (FP). The following were used to assess the performance of Missense3D-PPI: sensitivity, specificity, true positive rate (TPR), false positive rate (FPR), TPR/FPR ratio, accuracy and Matthews Correlation Coefficient (MCC).

Missense3D-PPI was benchmarked against Missense3D [33], MutaBind2 [17], BeAtMuSiC [16] and mCSM-PPI2 [18]. The last three methods are based on energy calculation and provide a ΔΔG value as an output. We defined variants as damaging if resulting in ΔΔG ≥1.5 kcal/mol or ≤ −1.5 kcal/mol, otherwise neutral [34]. McNemar’s test [35,36] was used to compare the performance of Missense3D-PPI against that of other algorithms.

## RESULTS

Missense3D-PPI was trained on a dataset of 640 variants (320 pathogenic and 320 benign; 435 occurring in a rim residue, 196 in a core and 9 in a support residue (Table S2). These variants were harboured by 310 human proteins and mapped onto 375 human protein complexes. On the training set, salt bridge and H-bond breakage/formation and charge-related features performed best on core and support residues. These features were, therefore, not used on rim residues in the testing set.

The performance of Missense3D-PPI was assessed on a test set, which comprised of 639 interface variants (413 damaging and 226 benign) occurring in 170 protein complexes and 124 unique human proteins. The 20 structural features analysed by Missense3D-PPI are presented in Table S1. In the test set, no variants caused a breakage of disulphide bonds, a feature that on the training set had obtained a TPR/FPR ratio of 7.4. All structural features showed a TPR/FPR >1, suggesting that Missense3D-PPI can accurately distinguish between damaging and neutral variants. Missense3D-PPI achieved an accuracy of 58% with a sensitivity (TPR) of 44% and a specificity of 83%. Missense3D-PPI performed better on core rather than rim residues (accuracy: 62% versus 56%, sensitivity: 63% versus 25%, specificity: 56% versus 91%. Only 16 residues were annotated as ‘support’, a number too small to allow us to calculate the accuracy of the predictor on this single type of interface residues.

The performance of Missense3D-PPI was compared to that of our in-house Missense3D, which assesses the effect of missense variants using the 3D coordinates of the monomeric protein. The performance of Missense3D-PPI was superior to that of Missense3D: sensitivity improved from 8% to 44% and the overall accuracy from 40% to 58%, p=4.23×10^−16^ (McNemar’s test). When comparing the accuracy of Missense3D-PPI to that of other structure-based predictors for variants at protein interface, Missense3D-PPI was consistently superior to BeAtMuSiC (p=2.3×10^−5^) and mCSM-PPI2 (p=3.2×10^−12^) and MutaBind2 (p=0.003, Table 1).

**Table 1.** Missense3D-PPI performance compared to other algorithms.

|  | Missense3D-PPI | Missense3D | MutaBind2 | BeatMusic | mCSM-PPI2 |
| --- | --- | --- | --- | --- | --- |
| **MCC** | 0.28 | 0.12 | 0.21 | 0.20 | 0.16 |
| **Accuracy** | 58% | 40% | 50% | 50% | 43% |
| **Sensitivity** | 44% | 8.5% | 27% | 27% | 14% |
| **Specificity** | 84% | 98% | 90% | 90% | 96% |
| **TPR** | 0.44 | 0.08 | 0.27 | 0.27 | 0.14 |
| **FPR** | 0.16 | 0.02 | 0.10 | 1.10 | 0.04 |

### Case study

Variant p.Ser336Arg causes medium-chain specific acyl-CoA dehydrogenase (ACADM) deficiency (OMIM Disease ID, <u>MIM: 201450</u>) [37]. This variant is harboured by the human enzyme ACADM (<u>UniProt ID P11310</u>). Serine 336 is located on the surface of the ACADM protein and its substitution to the large and charged arginine is predicted tolerated when assessed using the ACADM single chain 3D coordinates in Missense3D because it does not affect the correct folding and stability of the single protein. However, the same variant is predicted damaging by Missense3D-PPI when its effect is assessed on the ACADM homotetramer 3D structure (<u>PDB:4p13</u>; Figure 2 top panel). The substitution to arginine introduces an unbalanced charge at the protein interface. Furthermore, it is predicted to cause a steric hindrance of ACADM homotetramer assembly.

**Figure 2.**
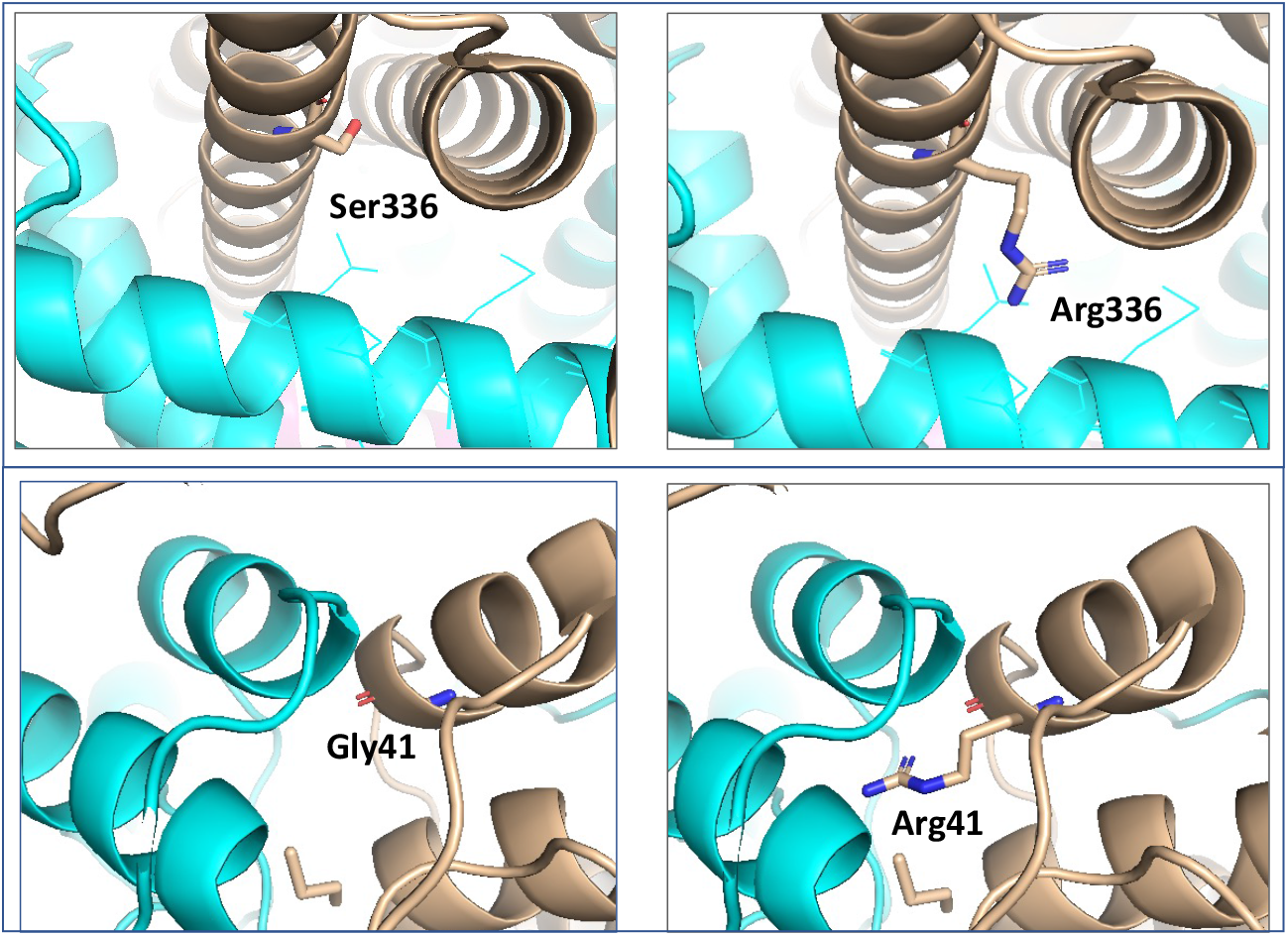
Damaging variants affecting protein interfaces are correctly identified by Missense3D-PPI: two case studies. Top panel, p.Ser336Arg in the human enzyme ACADM (PDB: 4p13). Bottom panel, p.Gly41Arg in the AGXT human enzyme (PDB: 4kyo).

Another example of the added value of assessing the effect of genetic variants with Missense3D-PPI is p.Gly41Arg in the alanine glyoxylate aminotransferase (AGXT) human enzyme (<u>UniProt ID P21549</u>), which causes type I primary hyperoxaluria (<u>MIM 259900</u>). Missense3D-PPI predicts this variant to be damaging through the formation of a new interchain hydrogen bond, the replacement of an interface glycine with any other residue and the introduction of an unbalanced charge at the AGXT homodimeric interface (<u>PDB:4kyo</u>; Figure 2, bottom panel). When analyzed on the single unbound AGXT structure, this variant is predicted tolerated by Missense3D because this amino acid substitution on the protein surface does not affect the AGXT monomeric protein structure.

### Web server

Missense3D-PPI is freely available at: (http://missense3d.bc.ic.ac.uk/missense3d/indexppi.html). Two Input pages are available to the user: if an experimental structure covering the residue harbouring the variant is available from PDB, the ‘Position on Protein Sequence” can be used. In this case the protein UniProt identifier and the position of the residue according to UniProt numbering should be provided. If a 3D model (including AlphaFold) or an experimental structure not yet deposited in PDB are used for the analysis of the variant, the “Position on 3D Structure” should be selected. In this case, a 3D coordinate file should be uploaded and the position of the variant specified according to the 3D structure residue numbering. Both Input pages require knowledge of the chain ID which identified the query protein in the 3D coordinates file. Example submissions entries are provided.

The initial output of Missense3D-PPI informs the user whether a structural damage has been identified or not. The in-depth structural analysis is provided in a separate Results page (Figure 3). The latter allows to visualize and manipulate the wild type and mutant residue on the 3D structure using a JSmol [38] interactive window. Furthermore, the structural damage, if any, identified is described along with all other structural features analysed.

**Figure 3.**
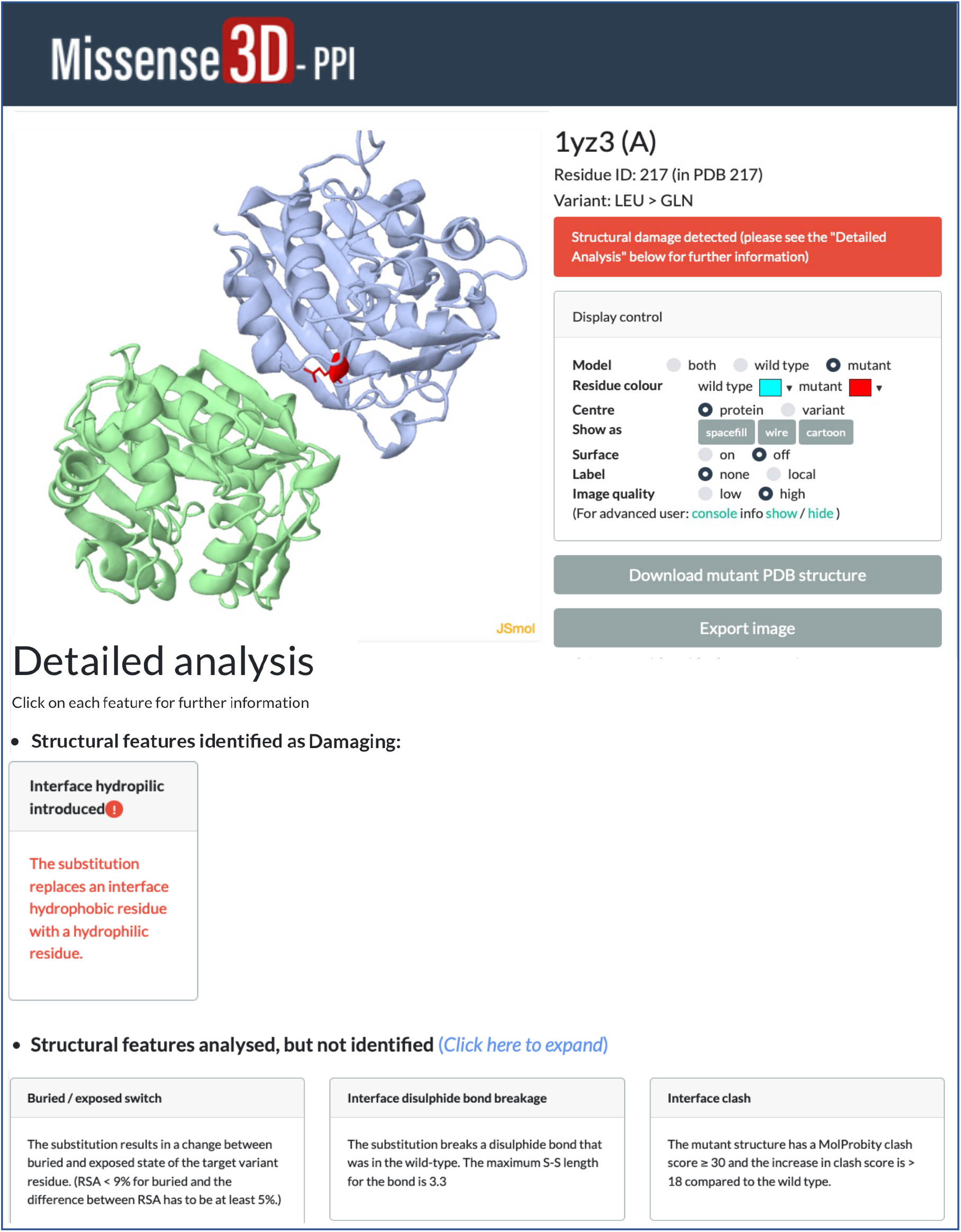
Missense3D-PPI Result page.

## DISCUSSION

We have developed Missense3D-PPI to specifically address the problem of predicting the effect of missense variants occurring on the surface of a protein in a PPI site. Missense3D-PPI requires the 3D coordinates of a protein complex, experimentally determined or modelled through algorithms, such as AlphaFold [12] or GWIDD [39], the latter available from the GWYRE database [9]. Moreover, additional precomputed 3D models of protein complexes are available from databases, such as Interactome3D [8] and PrePPI [10]. We showed that, for variants located at protein interfaces, Missense3D-PPI performed superiorly compared to Missense3D and to other structure-based variant predictors.

Missense3D-PPI is specifically designed to address the disruption of structural features, such as cavities, clashes or chemical bonds occurring in interchain residues. Such features are not included in our previous algorithm Missense3D, which uses the 3D coordinates of the monomeric protein. Another major difference between Missense3D-PPI and Missense3D is the introduction of interface-specific features, such as replacement of tyrosine and tryptophan, which are important residues for protein-protein interaction [31]. To account for these residues being enriched in interaction hot spots, we introduced the qualitative features “Interface Trp replaced” and “Interface Tyr replaced”, when either a tryptophan or a tyrosine interface residue is replaced by any other amino acid. Arginine has also been shown to contribute to PPI [31]. However, the replacement of arginine is already captured by features, such as ‘charge replaced’, ‘charge switch’ and ‘salt bridge breakage’, therefore an ‘interface arginine replaced’ feature was not introduced in Missense3D-PPI.

Two additional features, ‘interface Gly replaced’ and ‘interface Pro introduced’, were added in Missense3D-PPI to address potential structural problems which may occur at a protein interface. Indeed, glycine and proline, have unique structural features and their replacement/introduction can affect the integrity of a protein interface. Glycine is the smallest residue and its substitution with larger residues may introduce a clash, whereas introduction of a proline may limit the degree of flexibility of the backbone, thus affecting the conformation of the secondary structure of a protein interface [1] [40].

On our test set, “interface charge replaced”, “interface salt bridge breakage”, “interface Gly replaced”, “interface Tyr replaced” and “interface Gly in a bend” had the best TPR/FPR ratio, which is in line with the known physico-chemical properties of protein interfaces. “Breakage of a disulphide bond” was one of the top-performing features in the training test, but, unfortunately, no variant exhibited this feature in the test set. Disulphide bonds play an important role in protein structure stabilization and dimer formation [41] [42]. For this reason, the feature “disruption of disulphide bond” was kept in the final Missense3D-PPI algorithm.

Protein interfaces are not homogenous regions, and the properties of interface residues may vary according to their role within the interface, e.g. core or rim, and the type of protein interaction, e.g. transient or permanent. In our dataset, features, such as replacement of charged residues and disruption/formation of chemical bonds, only performed well when calculated on core residues. Indeed, core residues are more important in stabilizing the PPI compared to rim residues. Moreover, the performance of Missense3D-PPI should ideally be tested according to the nature of the protein interaction but, to our knowledge, no database of transient and permanent PPI is currently available. Homodimers have often been reported as nearly always permanent PPI, whereas hetero complexes as often transient, yet we did not identify a difference in the performance of Missense3D-PPI in homodimers versus heterocomplexes (data not shown).

Missense3D-PPI was benchmarked against three widely used energy-based programs for interface variants, BeAtMuSiC [16], MutaBind2 [17] and mCSM-PPI2 [18] and was significantly more accurate that all three. The added value of Missense3D-PPI compared to other programs includes a detailed description of the structural problem identified and the visualization of the mutant residue on the 3D structure. Moreover, in Missense3D-PPI the user does not need to specify the interacting chain in the 3D coordinate file, since the algorithm automatically calculates all interface residues between the protein of interest and other partners in the 3D coordinates file. This is particularly useful when the 3D coordinates file is that of a multimer.

A limitation of Missense3D-PPI is that predictions are solely based on the disruption of structural features and do not include information derived from sequence conservation. We therefore recommend using Missense3D-PPI as an additional tool to variant prediction, as recommended by the American College of Medical Genetics and Genomics (ACMG) and the UK Association for Clinical Genomic Science (ACGS) Best Practice guidelines on variant classification [43] [44]. Another limitation is that Missense3D-PPI requires the 3D coordinates of protein complexes. Although the coverage of the human proteome interactome is still quite limited, one can expect a substantial expansion in the 3D coordinates of protein complexes in the near future from development such as AlphaFold [12]. We did not test Missense3D-PPI on models, but, because it was developed using the same strategy adopted in Missense3D, we do not expect a significant drop in performance when applied to 3D models of complexes. In conclusion, Missense3D-PPI is a new resource to aid in the prioritization and characterization of genetic variants disrupting protein networks.

---

**Table S1.** Structural features analysed by Missense3D-PPI for residues participating in protein-protein interactions (interface residues).

| Features | Description | Interface residue |
| --- | --- | --- |
| Interface H-bond breakage/formation | The substitution breaks or forms side-chain/side-chain H-bond(s) and/or side-chain/main-chain bond(s) between two interchain residues. The maximum H-bond N–O length is 3.9Å. | Core, Support |
| Interface salt bridge breakage/formation | The substitution breaks or forms a salt bridge between two interchain residues. The maximum N–O bond length is 5.0Å. | Core, Support |
| Interface charge switched | The substitution replaces a charged interface residue with a residue of opposite charge [e.g. from arginine to aspartic acid]. | Core, Support |
| Interface charge introduced | The substitution replaces a non-charged interface residue with a charged one (histidine, lysine, arginine, glutamic acid or aspartic acid). | Core, Support |
| Interface charge replaced | The substitution replaces a charged interface residue (histidine, lysine, arginine, glutamic acid or aspartic acid) with a non-charged one. | Core, Support |
| Interface buried/exposed switch | The substitution results in a change between buried and exposed state of the target residue. (RSA < 9% for buried and the difference between RSA has to be at least 5%) | Core, Support |
| Interface hydrophilic introduced | The substitution replaces a hydrophobic interface residue with a hydrophilic one. | Core, Support, Rim |
| Interface disulfide bond breakage | The substitution breaks an interchain disulfide bond that was present in the wild-type structure. The maximum S–S length for the bond is 3.3 Å. | Core, Support, Rim |
| Interface clash | The mutant structure has a MolProbity clash score $\geq 30$ and the increase in clash score is $>18$ compared to the wild type. | Core, Support, Rim |
| Interface secondary structure altered | The substitution results in a change in the DSSP secondary structure assignment. | Core, Support, Rim |
| Interface cavity altered | The substitution leads to an expansion ($\geq 70 \text{ \AA}^3$) or contraction ($< 70 \text{ \AA}^3$) of the cavity volume. | Core, Support, Rim |
| Interface cis Pro replaced | The substitution replaces an interface proline, which was in cis configuration in the wild type. | Core, Support, Rim |
| Interface Gly in a bend | The substitution replaces an interface glycine, which is located in a bend curvature (reported “S” in DSSP). | Core, Support, Rim |
| Interface disallowed phi/psi | The mutant residue is in an outlier region, while the wild-type residue is in the favoured/allowed region. | Core, Support, Rim |
| Interface Pro introduced | The substitution introduces a proline at the interface. | Core, Support, Rim |
| Interface Gly/Tyr/Trp replaced | The substitution replaces an interface glycine, tyrosine, or tryptophan residue with any other residue. | Core, Support, Rim |

**Table S2.** Training and testing sets.

|  | Training set | Testing set |
| --- | --- | --- |
| Total variants | 640 | 639 |
| Pathogenic variants | 320 | 413 |
| Benign variants | 320 | 226 |
| Rim variants | 435 | 381 |
| Core variants | 196 | 242 |
| Support variants | 9 | 16 |
| Number of proteins | 310 | 124 |
| Number of complexes | 375 | 170 |

